# Improving wearable-based seizure prediction by feature fusion using growing network

**DOI:** 10.1101/2025.01.28.635212

**Authors:** Tanuj Hasija, Maurice Kuschel, Michele Jackson, Stephanie Dailey, Henric Menne, Claus Reinsberger, Solveig Vieluf, Tobias Loddenkemper

**Author notes:** Shared first authorship (Tanuj Hasija, Maurice Kuschel). Shared last authorship (Solveig Vieluf, Tobias Loddenkemper). { }. {, }.

## Abstract

The unpredictability of seizures is one of the most compromising features reported by people with epilepsy. Non-stigmatizing and easy-to-use wearable devices may provide information to predict seizures based on physiological data. We propose a patient-agnostic seizure prediction method that identifies group-level patterns across data from multiple patients. We employ supervised long-short-term networks (LSTMs) and add unsupervised deep canonically correlated autoencoders (DCCAE) and 24-hour patterns using time-of-day information. We fuse features from these three techniques using a growing neural network, allowing incremental learning. Our method with all three features improves prediction accuracy over the baseline LSTM by 7.3%, from 74.4% to 81.7%, averaged across all patients, and outperforms the LSTM in 84% of patients. Compared to the all-at-once fusion, the growing network improves the accuracy by 9.5%. We analyze the impact of preictal data duration, wearable data quality, and clinical variables on the prediction performance.

## 1 Introduction

Epilepsy is a neurological condition that affects more than 50 million people in the world [1]. It ranks fourth on the world’s disease burden list with a worldwide yearly healthcare burden estimated at 110 billion euros [2]. In children, epilepsy is the most frequent chronic pediatric neurological condition. Epilepsy presents with abnormal electrographic brain activity resulting in often debilitating seizures, which can cause unconsciousness, physical injuries, cognitive disability, neurological injury, as well as stressors and reduced quality of life for patients and families alike. Further, seizures can lead to sudden death in epilepsy (SUDEP) with an incidence rate of approximately 1 in 1,000 people with epilepsy each year [3, 4]. Moreover, generalized tonic-clonic (GTC) seizures, one of the most debilitating seizure types, are the greatest risk factor for SUDEP with rates as high as 1 in 100 for those experiencing frequent seizures [5]. One-third of people with epilepsy continue to experience recurrent seizures despite treatment [6].

The unpredictability of seizures, i.e., not knowing when the next seizure will occur, is one of the most compromising features reported by people with epilepsy and their caregivers [7]. Even though a rescue medication to prevent or abort a seizure exists in many cases and reduces the likelihood of SUDEP, the unpredictability of an impending seizure leads to uncertainty regarding when to take the rescue medication [8]. Therefore, an effective seizure prediction algorithm that can provide an accurate likelihood of an impending seizure has the potential to save lives and improve the quality of life for people with epilepsy.

Most seizure prediction methods are based on electroencephalography (EEG) [9, 10, 11] and seizure diaries [12], with varying performance between patients and seizure types [13, 14]. One study demonstrated the viability of seizure prediction using intracranial EEG devices implanted in the brain [9]. However, this study also revealed that approximately two-thirds of individuals experienced side effects from the implanted devices. Therefore, such devices may not be appropriate for widespread use in the general population, especially in children, as they can interfere with normal brain development. Seizure prediction algorithms using noninvasive scalp EEG devices have been developed using various methods ranging from feature extraction-based methods to completely data-driven deep learning techniques [10, 11]. However, such devices are associated with high costs and carry a stigma due to the numerous electrodes placed on the scalp [15].

Non-stigmatizing, low-cost, and easy-to-use wearable devices such as wristbands deployed to the wrist and ankle that collect autonomic nervous system (ANS) data from modalities such as heart rate (HR), electrodermal activity (EDA), and skin temperature (TEMP), provide a potential alternative to EEG [16]. These wearable devices are equipped with a variety of sensors, such as accelerometer (ACC), photoplethysmography (PPG), electrocardiogram (ECG), and EDA sensor, which have also been employed in many commercial smartwatches [17]. Thus, wearables offer a feasible option for seizure prediction in an outpatient setting, especially for children.

One pre-requisite for an effective seizure prediction algorithm is to differentiate between the data recorded during the interictal period (not close to seizures) and data recorded during the preictal period (shortly before a seizure) with high accuracy. Research on seizure prediction has shown that seizures are associated with preictal changes in the ANS modalities, which contain information for predicting seizures [18, 19, 20, 21]. Seizure prediction using wearables is an emerging field, supported by recent proof-of-concept techniques demonstrating the potential of these methods [22, 23, 24, 25, 26].

Most of these techniques are based on supervised learning, where labeled preictal and interictal class information is paired with time-series data or features [27]. For instance, in [22], a naive Bayes classifier was trained using the mean values of EDA, HR, TEMP, and blood volume pulse (BVP) measured from the PPG sensor as features. In [23], the authors trained a long short-term memory (LSTM) neural network on filtered time-series data from EDA, ACC, BVP, and TEMP, while [26] involved training an LSTM on raw data from EDA, ACC, BVP, TEMP, HR, along with their Fourier-transformed data, data-quality features, and time-of-day features. In [24], three different supervised learning methods were employed: LSTMs trained on daily sleep features, a random forest (RF) regressor trained on cyclic, HR, and physical activity features, and a logistic regression ensemble combining the outputs of the LSTM and RF models to predict the final seizure risk. Conversely, unsupervised learning techniques obtain features from data to optimize a specific cost function, such as maximizing the variance of the features rather than using class label information. For example, in [25], the authors extracted heart rate variability (HRV) data from a custom ECG wearable device and applied anomaly detection using principal component analysis (PCA) and statistical indices to control the false alarm rate.

An important aspect often overlooked in the techniques mentioned above is the complex interconnection of subsystems that form the ANS. The state-specific interplays of several organ-specific subnetworks are crucial for maintaining the functionality of the ANS, which operates through centrally coupled and modality-specific control mechanisms [28]. The changes in interactions among these ANS subsystems preceding an epileptic seizure present a promising avenue for seizure prediction [21]. Several studies have shown that impending seizures could alter the central control of the autonomic network, causing coupled changes in the ANS subsystems with distinct temporal patterns [29, 30, 31]. In this context, canonical correlation analysis (CCA) provides an effective, data-driven method for measuring multimodal interactions among different modalities [32]. CCA is an unsupervised learning method that identifies features from each modality that are maximally correlated. This method extracts common (or highly correlated) features across modalities while discarding noisy (or uncorrelated) data from individual modalities. In [31], a combination of PCA and CCA reliably detected preictal coupled changes across EDA, HR, TEMP, and respiratory rate (RR). However, a significant limitation of CCA is its capacity to extract only linear relationships between multimodal data [33]. Given the complexity and nonlinearity of seizure-induced changes in the ANS, it is reasonable to assume that these relationships are highly complex and nonlinear [34]. Deep CCA (DCCA) extracts highly correlated features from multimodal data with nonlinear relationships [35]. Deep canonically correlated autoencoders (DCCAE) improve DCCA by addressing potential overfitting using autoencoder regularization in conjunction with DCCA [36]. A DCCAE-based method has recently demonstrated the feasibility of a seizure prediction biomarker by classifying preictal data from seizure patients and interictal data from patients without seizures [37].

In addition to the autonomic data, epileptic seizures exhibit 24-hour and multi-day patterns, with seizure likelihood varying based on the time of day [38, 39, 40]. In [41], probability density functions are used to approximate seizure-specific cyclic patterns, and incorporating this improved the prediction performance of a baseline logistic regression model applied to electrocorticography data. [42] integrated wearable patient-specific data and time-of-day information using LSTM and reported that the time-of-day features contributed the most to the area under the curve (AUC). Therefore, features derived from 24-hour patterns add complementary information and could enhance the accuracy of seizure prediction models.

In this study, we hypothesize that different features focus on different types of biomarkers and include complementary information, and their combination could improve prediction accuracy. Our contributions to the existing work are: i) We propose a novel fusion of complementary features for wearable-based seizure prediction: supervised features using LSTM, canonical correlation-based multimodal features using DCCAE, and 24-hour patterns using time-of-day features. ii) We employ a growing neural network to integrate different features and incrementally expand the fusion network size to include additional features. This approach is evaluated with the typically used all-at-once feature fusion approach. iii) We analyze how the quality of data from different autonomic modalities, clinical variables, and preictal data duration affects prediction performance.

## 2 Materials and methods

### 2.1 Dataset

#### 2.1.1 Ethics statement and patient consent

We conducted this study following the Institutional Review Board (IRB) guidelines and regulations at Boston Children’s Hospital, applicable government regulations, and the Declaration of Helsinki. Boston Children’s Hospital IRB approved the study (Autonomic biomarkers of seizures to assess the risk for SUDEP, IRB-P00001526). We obtained written informed consent and assent from all participants and/or their guardians before enrollment.

#### 2.1.2 Patient selection and clinical data inclusion

We included patients who were enrolled in the long-term monitoring (LTM) unit with video-EEG at Boston Children’s Hospital between February 2015 and February 2021 and wore a wearable sensor (Empatica® E4, Milan, Italy) on their wrist and/or ankle that recorded EDA, TEMP and HR. During this period, a total of 450 patients were enrolled. A board-certified epileptologist (TL) scored seizures and determined electroencephalographic seizure onset times. The inclusion criteria are summarized in Fig. 1.

**Figure 1:**
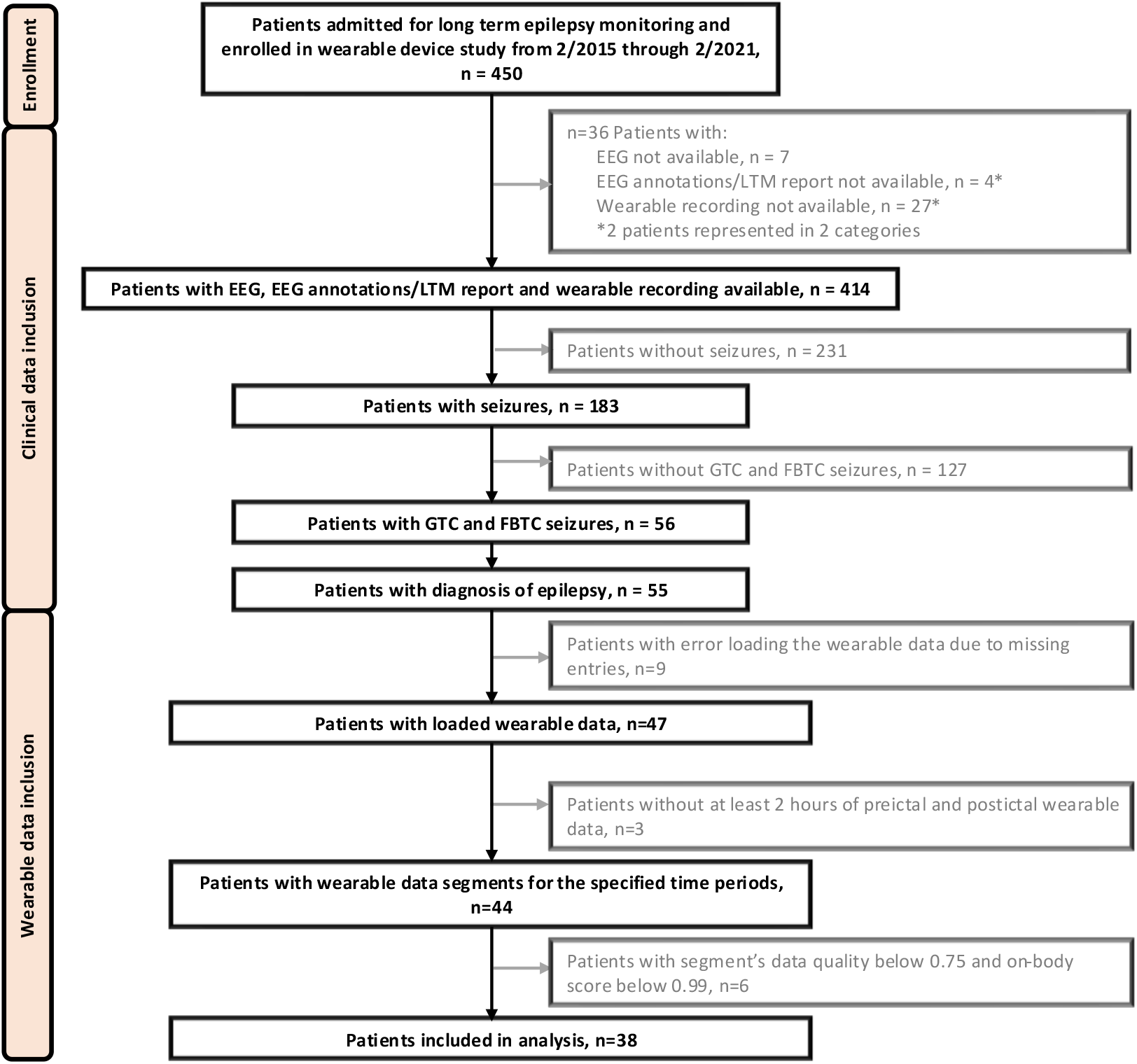
The inclusion diagram shows the patient selection criteria based on clinical data and wearable data. Of the 450 enrolled patients, we excluded patients with unavailable EEG data, LTM reports, and wearable recordings and those with no seizures during their LTM stay. From the remaining 183 patients, we included patients with confirmed epilepsy diagnosis and those who had tonic-clonic, i.e., GTC and focal to bilateral tonic-clonic (FBTC) seizures.

#### 2.1.3 Wearable data inclusion

We further excluded patients with missing data entries (Fig. 1). The remaining patient data was segmented into five different segments with the following criteria:

a. Preictal segment: Two hours of data preceding the 30 seconds immediately before seizure onset. We chose 30 seconds as a buffer for synchronization inaccuracies due to clock drift between the wearable and EEG devices [43].
b. Postictal segment: Two hours of data following the 30 seconds after seizure offset.
c. Interictal segment: Data at least two hours and 30 seconds away from seizure onset and offset.
d. Ictal segment: Data between seizure onset and offset, including the 30-second buffer at both ends.
e. Invalid segment: Data that does not satisfy either a), b), c), or d). For example, the segment between two seizures less than two hours apart or seizure immediately (less than two hours) after patient enrollment.

Fig. 2 shows an example of the labeled segments from one patient’s enrollment data. Our criteria exclude seizures less than 4 hours and one minute apart. Only patients with at least one valid preictal and interictal segment are used for further analysis (Fig. 1).

**Figure 2:**
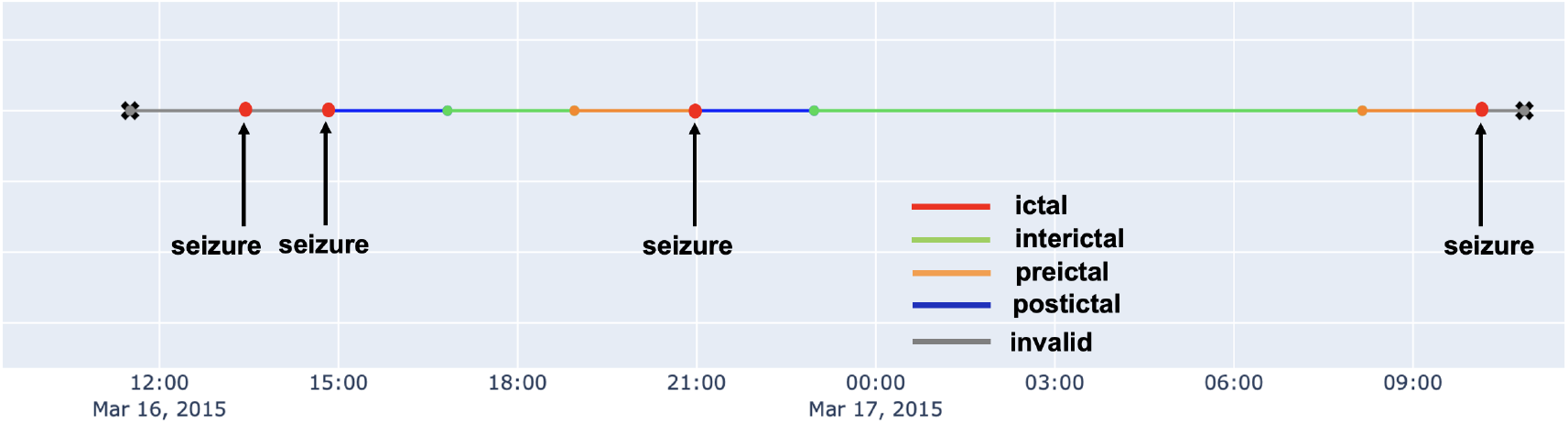
Data and labeling showing five segments with different colors for one patient’s enrollment data. The ictal segment is shorter than the other segments and is only visible as a red dot in the figure.

#### 2.1.4 Data quality assessment

To assess the reliability of each preictal and interictal segment, we compute the onbody score and the data quality as follows.

i. Onbody score: The onbody score indicates how long the wearable was worn on the body during data recording [44]. It takes into account ACC, TEMP, and EDA data. Every modality’s data is compared against a pre-defined threshold as proposed in [44]. The onbody score of a segment is the average of all modalities’ onbody scores and ranges from 0 to 1. In our experiments, we excluded segments with an onbody score below 0.99.
ii. Data quality score: The data quality indicates the quality of the recorded onbody data. It is individually computed on ACC, TEMP, EDA, and BVP data and is implemented in Python using the methods defined in [44]. The data quality score of a segment is the average of all modality’s data quality scores and ranges from 0 to 1. We only considered segments with a score above 0.75, ensuring that no modality’s data has a quality of 0. If the patient had multiple recordings from both the ankle and wrist, we considered segments from recordings with higher data quality. In total, 38 patients are included in the analysis, with a total of 59 seizures from them (Fig. 1).

### 2.2 Data pre-processing

The preictal and interictal segments are divided into non-overlapping windowed segments, each comprising 15 seconds. Data from HR is upsampled to 4 Hz using the Python function *interpolate* so that all modalities have the same sampling frequency [45].

## 3 Proposed method

This section presents the proposed method for fusing features from multiple ANS modalities using LSTM, DCCAE, and time-of-day features. The complete processing pipeline is shown in Fig. 3. The wearable data comprising EDA, HR, and TEMP is pre-processed and filtered for data quality as described in Section 2. The pre-processed EDA and HR data is then fed into the DCCAE, described in Section 3.2, which extracts highly correlated features between these two modalities. Concurrently, the EDA, HR, and TEMP data are inputs to the supervised LSTM, described in Section 3.1, to capture temporal dependencies and classification-specific features. To account for 24-hour influences, the time-of-day information is encoded using two distinct methods described in Section 3.3. The extracted feature vectors from the DCCAE, LSTM, and time-of-day encoding are combined using a growing neural network, described in Section 3.4. The trained network finally classifies each input segment as interictal or preictal, predicting an impending seizure.

**Figure 3:**
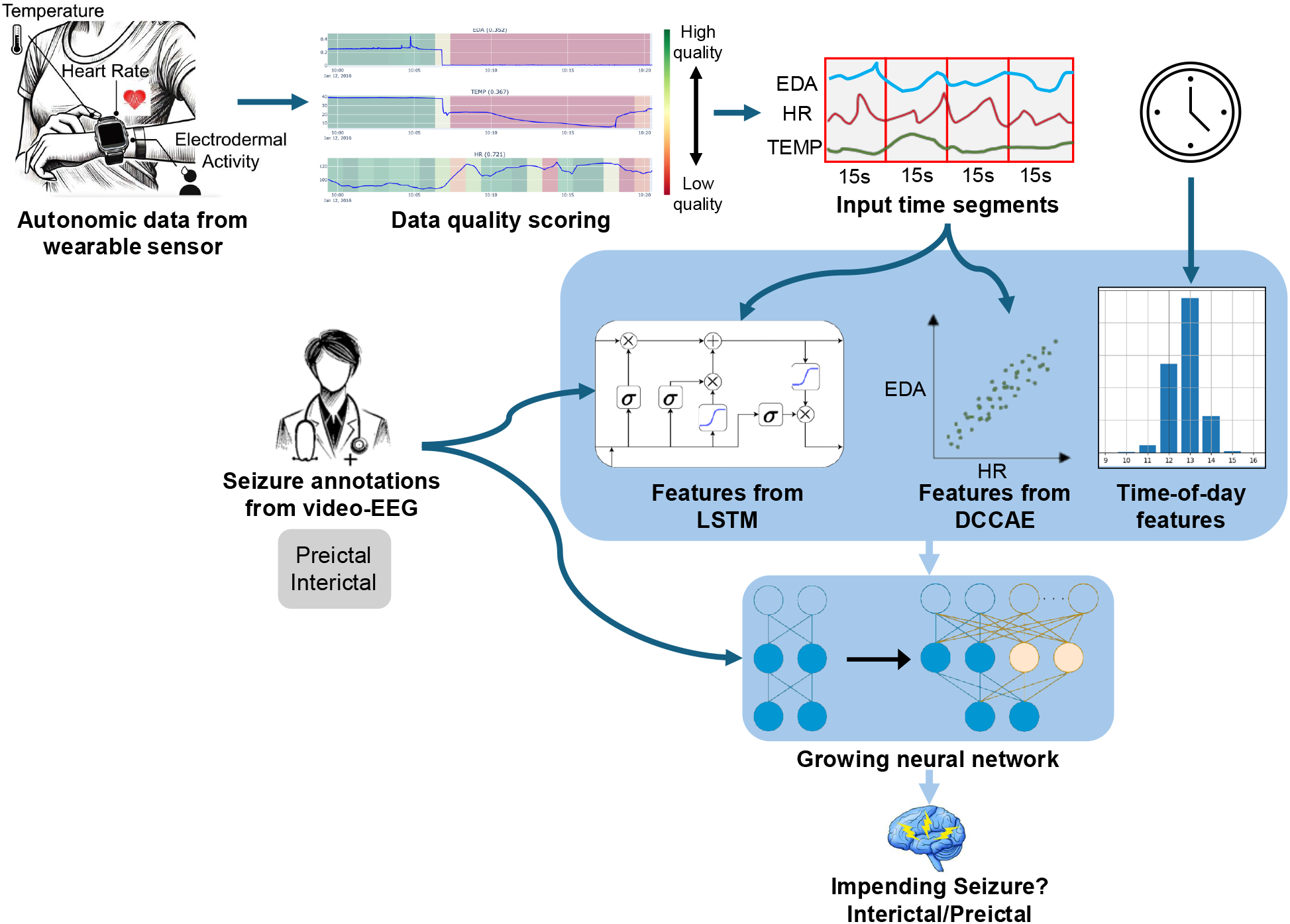
Proposed seizure prediction method. The ANS data from the wearable sensor comprising EDA, HR, and TEMP is pre-processed and filtered for data quality. The data is then segmented into windows of 15-second duration. The time segments are inputted into the supervised LSTM to capture temporal dependencies and classification-specific features, whereas the unsupervised DCCAE extracts highly correlated features using EDA and HR time segments. To account for 24-hour patterns, the time-of-day information is encoded using trigonometric and radial basis functions (RBF). The extracted feature vectors from the LSTM, DCCAE, and time-of-day encoding are combined using a growing neural network, which finally classifies each input segment as interictal or preictal, predicting an impending seizure.

### 3.1 Classification-specific features from supervised LSTM networks

An LSTM is a recurrent neural network (RNN) that extracts compact, low-dimensional features from time-series data [46]. In contrast to classic RNNs, it uses an LSTM cell designed to utilize long-term dependencies in the input sequence by comprising an internal state and different internal gates for the input and the output. When feeding a series of data into the cell, the final output is a low-dimensional representation of the input sequence. We present a schematic of the implemented LSTM cell in Fig. 4. We employ the LSTM for supervised learning to extract features from the wearable data that are specifically useful for classifying preictal and interictal segments. Specifically, we feed windows of 15 seconds from EDA, HR, and TEMP into the LSTM. Data of the modalities is stacked, resulting in an input shape of 60 × 3, as the data is sampled at 4 Hz. We set the dimensionality of the LSTM to be 15, resulting in the 15-dimensional output feature vector comprising low-dimensional features from all three modalities. The features extracted are fed to the growing neural network explained in Section 3.4.

**Figure 4:**
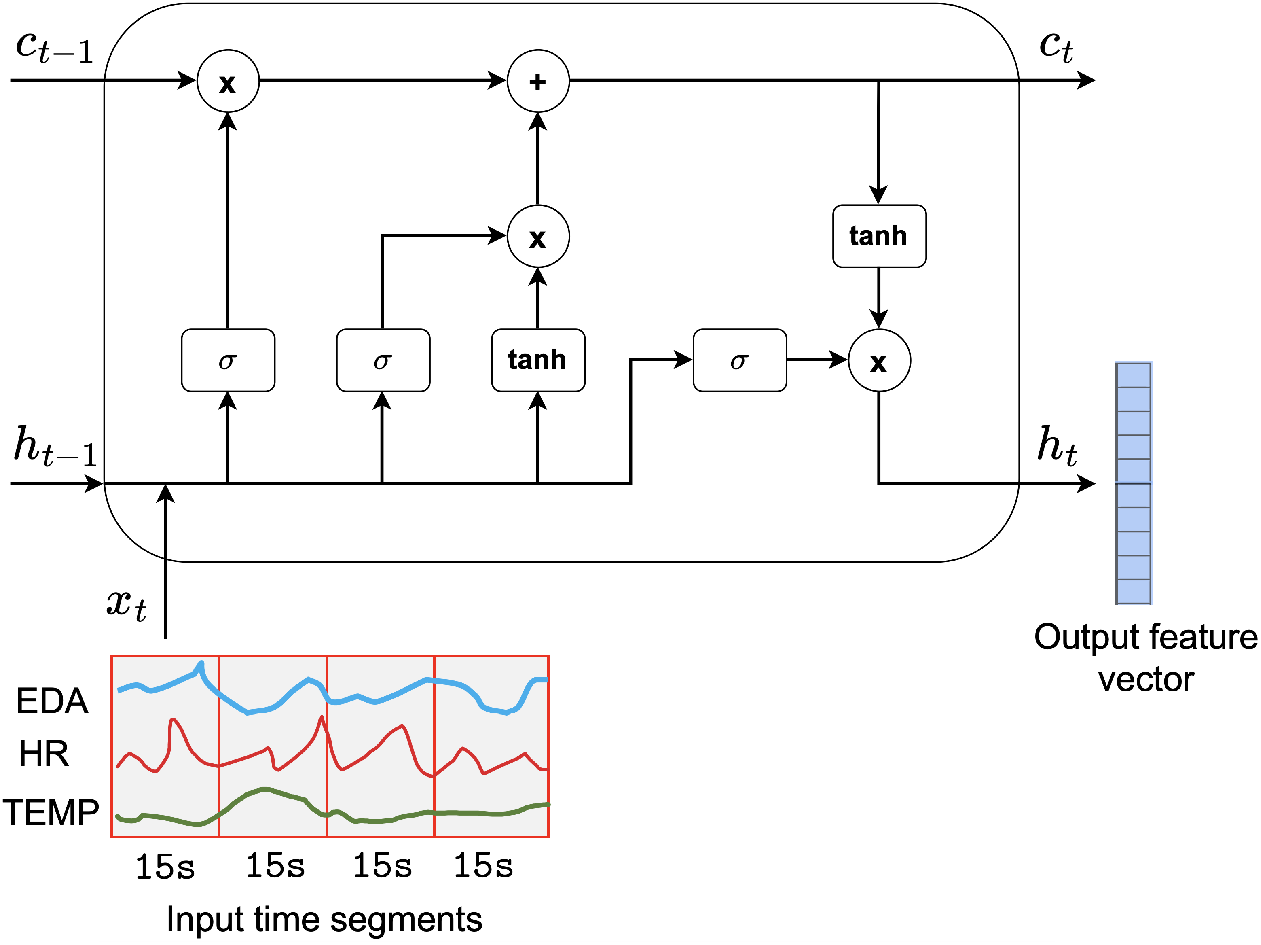
Schematic of the implemented supervised LSTM for extracting low-dimensional classification-specific features from EDA, HR and TEMP.

### 3.2 Correlation-based features from unsupervised DCCAE

DCCAE is an extension of traditional CCA for extracting nonlinear relationships between the data from two modalities [35, 36]. DCCAE transforms the input data to low-dimensional features using neural networks and maximizes the correlation between the transformed features of the two modalities. Let **E** ∈ ℝ^*t×N*^ denote the matrix containing all the preictal and interictal 15-second windows for EDA, and similarly, **H** ∈ ℝ^*t×N*^ denotes the matrix for HR. Here *t* is the number of time points in each window, and *N* is the total number of windows from all patients. Let **f** : ℝ^*t*^ → ℝ^*d*^, **g** : ℝ^*t*^ → ℝ^*d*^ denote the encoding nonlinear transformation functions implemented using LSTM neural networks and 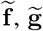 denote the decoding functions implemented using fully connected neural networks, as shown in Fig. 5. Here, *d* denotes the dimension of the low-dimensional features, and we chose *d* = 5. Further, **U** ∈ ℝ^*d×d*^ and **V** ∈ ℝ^*d×d*^ denote square matrices. DCCAE aims at learning 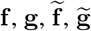 and **V** such that the correlation between the feature matrix of EDA, **P** = **U**^*T*^ **f** (**E**) and the feature matrix of HR, **Q** = **U**^*T*^ **f** (**E**) is maximized [36]. DCCAE employs an autoencoder regularization and certain constraints to avoid overfitting neural networks to trivial solutions. More specifically, the outputs of the encoders, **f** (**E**) and **g**(**H**) are also passed through the decoder neural networks, 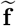 and, 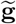 such that the reconstruction error between the decoder outputs and the original input data is minimized along with correlation maximization. Furthermore, each row of **P** and **Q** is constrained to be of unit variance, ensuring that trivial solutions with large values are avoided. Moreover, it constrains a one-to-one correlation between the features in **P** and **Q** and, thus, avoids redundant solutions. The optimization problem is

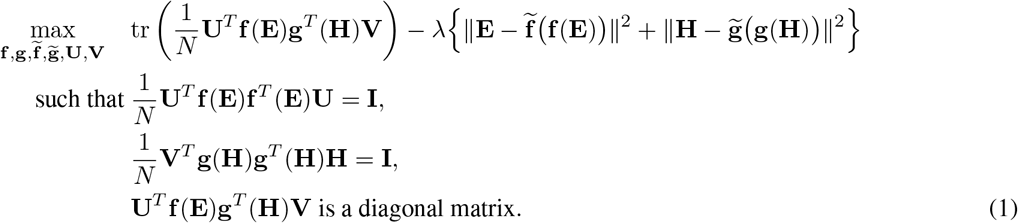

**Figure 5:**
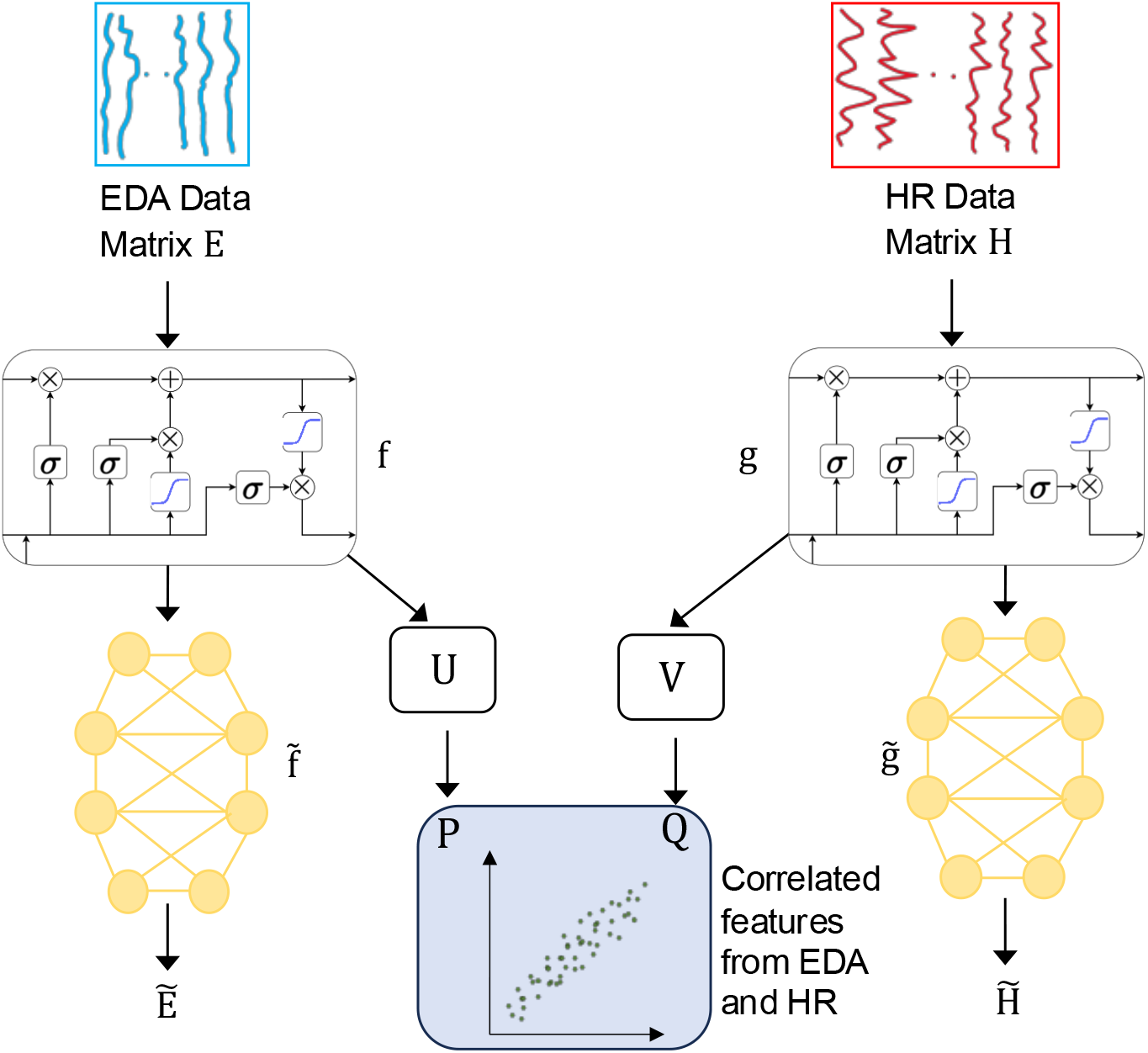
Unsupervised DCCAE for learning correlated features between the data from EDA and HR using correlation maximization and autoencoder regularization.

Here, ∥ · ∥ ^2^ denotes the norm and *λ* denotes the regularization parameter chosen as 1*e*^−7^ [36]. The feature matrices, **P** and **Q**, learned by DCCAE are inputted to the growing neural network explained in Section 3.4.

### 3.3 Time-of-day features

To include 24-hour seizure occurrence patterns in the proposed prediction method, we implemented two methods. These two methods encode the time of day using trigonometric functions and RBFs as follows.

#### 3.3.1 Trigonometric encoding

We use sine and cosine trigonometric functions to encode the time of day. A timestamp in the 24-hour format *hh* : *mm* : *ss* is first converted into *T* seconds. The sine and cosine values are then calculated based on the total duration of a day, which consists of 86, 400 seconds. The encoded values are given by

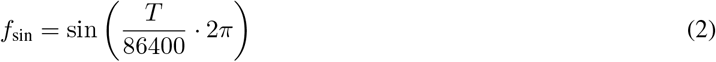

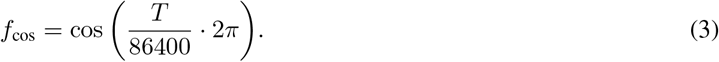

These two encodings are necessary because using only one of them, either sine or cosine, would result in ambiguity, as the same value could correspond to two different times of the day. Fig. 6a illustrates a time-of-day encoding using trigonometric functions.

**Figure 6:**
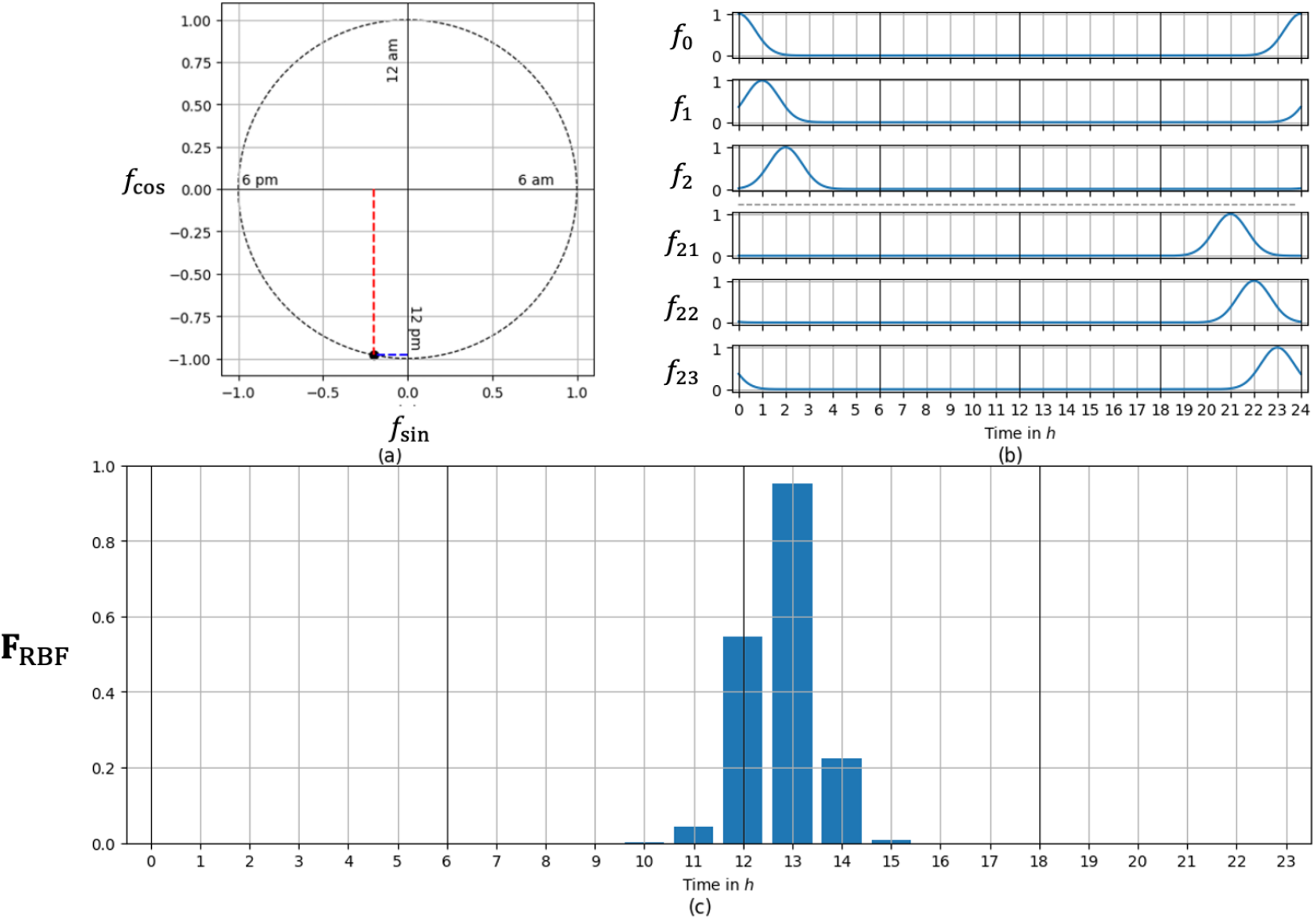
Example of time-of-day encodings for the timestamp 12:46:40 (a) the encoded value is denoted by the black dot using trigonometric functions with *f*_sin_ (red dashed line) and *f*_cos_ (blue dashed line) (b) RBFs corresponding to the first three and the last three hours of the day (c) the encoding using 24 RBFs.

#### 3.3.2 RBF encoding

An RBF is a function that represents the proximity to a specific point, often referred to as the center [47]. We generated the RBFs using the *RepeatingBasisFunction* from the scikit-lego package [48]. Each RBF is a Gaussian function, with the center represented by a mean value and a standard deviation of one. We selected 24 RBFs, *f*_0_, *f*_2_, …, *f*_23_, where each center corresponds to an hour of the day. Fig. 6b shows the RBFs for the first three and last three hours of the day, showing the cyclic pattern that reflects the repeating nature of time throughout the day. Each timestamp is encoded into 24 values,

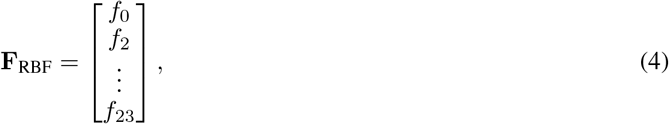

where all values are normalized between zero and one. A value of one indicates a perfect match between the timestamp and the center of an RBF, i.e., the timestamp matches exactly with a specific hour. In contrast, values close to zero indicate that the timestamp is far from the corresponding hour. Therefore, **F**_RBF_ can be seen as a “soft” one-hot encoding. An example of time-of-day encoding using RBFs is shown in Fig. 6c.

### 3.4 Growing neural network for fusing multiple feature vectors

For the final classification of each input segment, we fuse the extracted feature vectors from the supervised LSTM encoder, the unsupervised DCCAE, and the time-of-day encodings. Prior to fusing, we normalize all feature vectors to be in a comparable value range. LSTM and DCCAE features are normalized to be zero-mean and unit-variance, while features from both the time-of-day encodings are already normalized to be in [0, 1]. This prevents the final classification network’s training from focusing too much on individual feature vectors. The simplest option for the fusion of feature vectors is to create and train a new network from the beginning. However, the supervised LSTM encoder is already trained with a network specialized to classify segments based on the extracted features. Therefore, we utilize this existing network as a starting point for the combined network and grow its architecture during training. There are different approaches for incrementally growing a neural network [49, 50, 51], Here, we use a technique that expands the width of the neural network while preserving its output [52]. This has the advantage that the larger network immediately performs as well as the original network. Further, any local step in the training that is an improvement will be a guaranteed improvement over the original model. In practice, this ensures that the performance of the growing network will always be equal to or better than the baseline supervised LSTM alone. Such a guarantee does not exist for training a new network from the beginning or with other network-growing techniques.

### 3.5 Experimental Setup

The encoder of the supervised LSTM network consists of one 15-dimensional layer, followed by a layer for batch normalization. The extracted features are input to a fully connected layer with five hidden neurons, followed by a fully connected output layer with two neurons. L2 regularization is done with a regularization weight of 1*e*^−4^. The network is trained with categorical cross-entropy loss for 2000 epochs and evaluated on the evaluation split every five epochs. For the DCCAEs, the input dimension, *t*, is set to 60 and the encoder dimension, *d*, is set to 5 for both HR and EDA. The decoders are implemented as fully connected layers with linear activation and *t* = 60 units to match the input shape. We add a regularization term for computing the covariance matrices for CCA with the regularization parameter set to 1*e*^−4^ and L2 regularization on the encoders with the weight set to 1*e*^−4^. Training is done for 1000 epochs, and clustering accuracy is evaluated on the evaluation split every five epochs. The growing neural network is a fully connected neural network with two layers, similar to the supervised LSTM described before. However, it consists of 10 hidden neurons, where five neurons are taken from the supervised LSTM and then are duplicated. For the training of the growing network, the encoders of the DCCAEs and supervised LSTMs are fixed and not part of the optimization problem. Optimization in every experiment is done with Adam optimizer [53] and a learning rate of 1*e*^−3^. Training data is organized in one batch to allow for better extraction of group-level patterns. All experiments are run for ten different initializations.

### 3.6 Performance evaluation

To evaluate the performance of our method, we use the following metrics:

1. Accuracy: Measures the proportion of correctly classified preictal and interictal segments relative to the total number of segments.
2. Sensitivity: Measures the method’s ability to correctly classify preictal segments relative to the total number of preictal segments.
3. Specificity: Measures the ability to correctly classify interictal segments relative to the total interictal segments. High specificity ensures the method is prone to low false alarms.
4. AUC (Area Under the Curve): The AUC score represents the area under the ROC (Receiver Operating Characteristic) curve. The ROC curve plots the true positive rate against the false positive rate.

We evaluated our proposed method with 10-fold cross-validation, as shown in Fig. 7a, and reported the maximum accuracies on evaluation data, averaged over all folds. We used the evaluation data of each fold for early stopping to prevent the method from overfitting. For every fold, we compare the performance of all ten initializations and report the metrics belonging to the initialization with the best-performing model. To evaluate the proposed method on a patient level, we performed a leave-one-patient out cross-validation, as shown in Fig. 7b. For every patient, we excluded that patient’s data from the dataset. The remaining data is then subdivided into training and evaluation by an inner cross-validation. This type of evaluation gives insights on a patient level and also evaluates the generalization of the growing network’s performance improvement for unseen patients.

**Figure 7:**
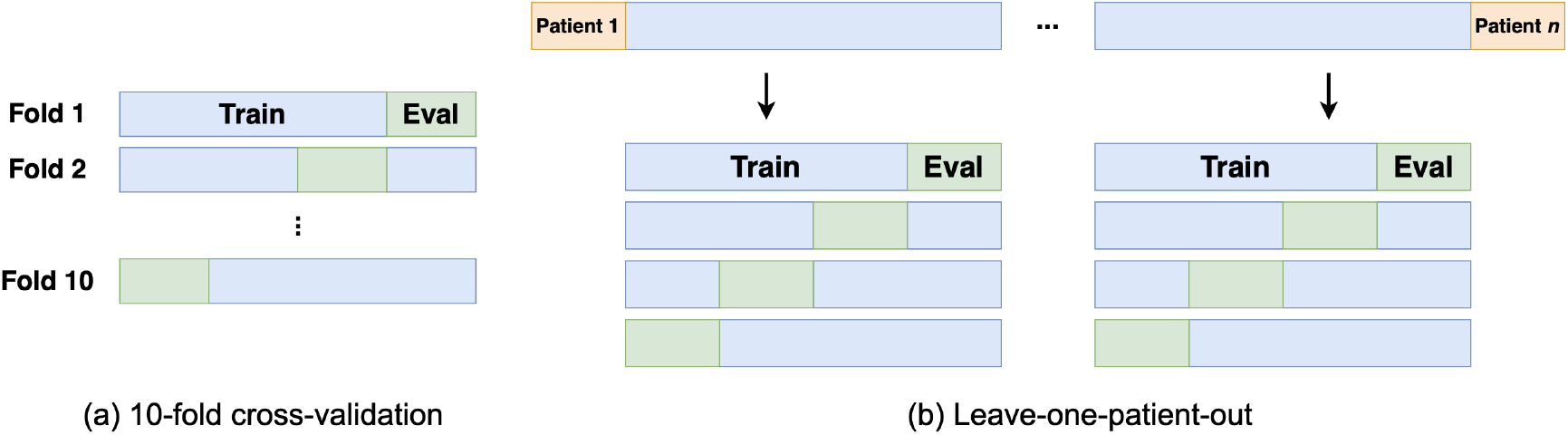
Visualization of the two evaluations used a) 10-fold cross-validation and b) leave-one-patient-out cross-validation.

## 4 Results

### 4.1 Dataset statistics

Fig. 8 illustrates the dataset statistics for the patients and seizures included in our analysis. Fig. 8a and 8b show the distribution of the two types of tonic-clonic seizures and the patients’ sex, respectively. Fig. 8c presents the histogram of the anti-seizure medications administered during the LTM stay in the hospital. Fig. 8d illustrates the distribution of etiology types among the patients, while Fig. 8e, 8f, and 8g display the histograms of age, the number of tonic-clonic seizures per patient, and the time of seizure onset, respectively.

**Figure 8:**
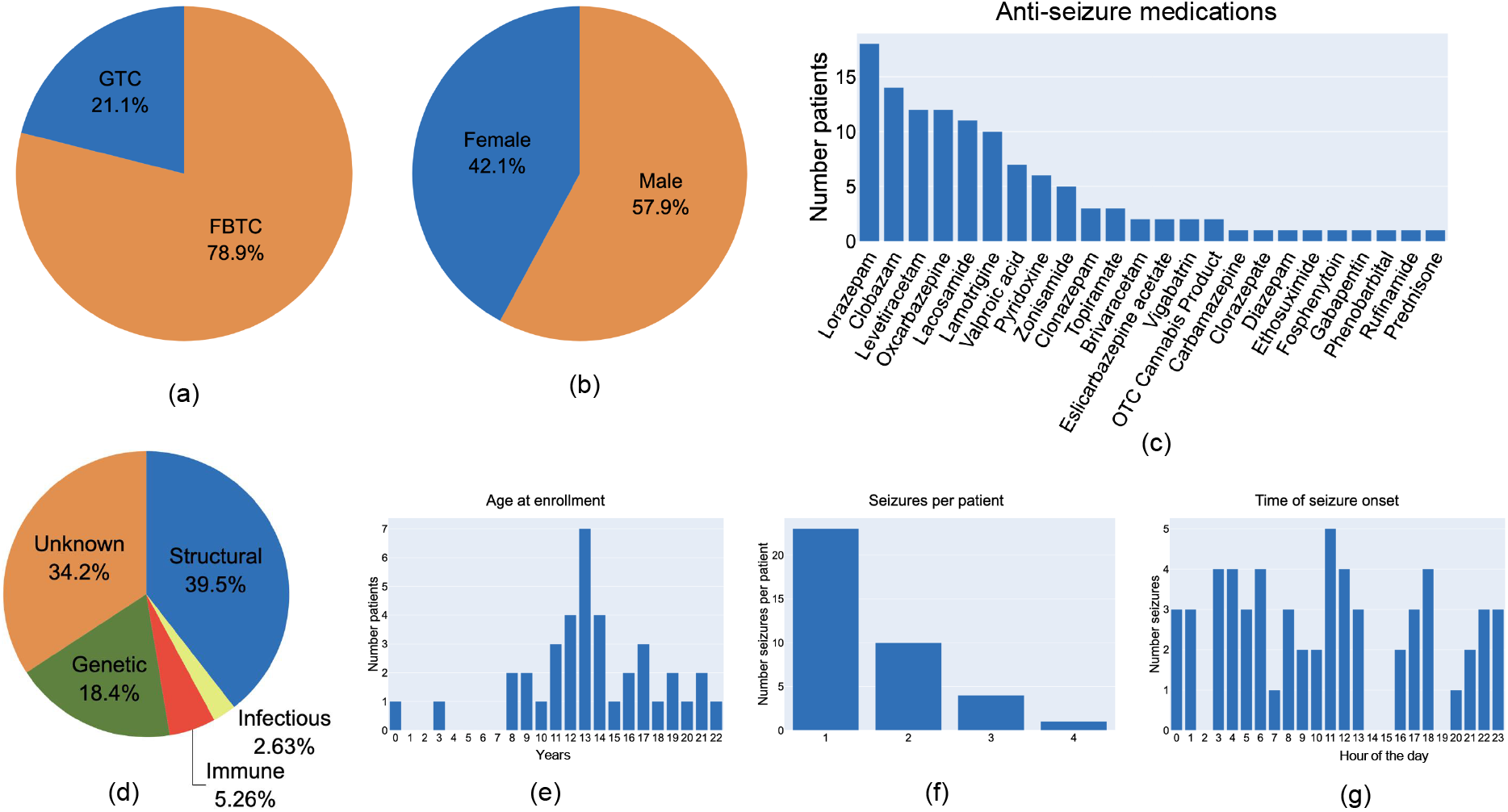
Dataset statistics of included patients and seizures: a) distribution of tonic-clonic seizure types, b) sex distribution, c) histogram of anti-seizure medications administered during LTM (patients may have been represented in more than one category), d) etiology distribution, histograms of e) age of patients at enrollment, f) number of seizures per patient during their LTM stay and g) time of seizure onset during the day.

### 4.2 Fusion of features from LSTM, DCCAE and time-of-day

The four performance metrics for the proposed method with 10-fold cross-validation are presented in Table 1. The results are averaged over all folds. We used the supervised LSTM technique as the baseline model as it achieved the best individual performance among the three different features. The LSTM alone achieves an accuracy of 74.44%, while the fusion of LSTM and DCCAE features using growing network provides a combined accuracy of 76.65%. Incorporating the time-of-day features in addition to the features from the LSTM and DCCAE improves the performance with the fusion of trigonometric encoding, providing 78.33% accuracy, and with the RBF encoding resulting in the highest accuracy of 81.70%. The fusion of the three feature sets with RBF encoding also results in the highest specificity and AUC.

**Table 1:** Evaluation of the proposed method fusing features from LSTM, DCCAE and the time-of-day using two different encodings for different performance metrics.

| Method | Accuracy | Sensitivity | Specificity | AUC |
| --- | --- | --- | --- | --- |
| LSTM | 74.44% | **82.40%** | 66.48% | 71.88% |
| LSTM + DCCAE | 76.65% | 77.84% | 75.45% | 72.75% |
| LSTM + DCCAE + Time-of-day (Trigonometric) | 78.33% | 82.09% | 74.58% | 76.58% |
| LSTM + DCCAE + Time-of-day (RBF) | **81.70%** | 79.34% | **84.07%** | **80.60%** |

The average ROC curves for the proposed method with RBF encoding and the baseline LSTM method are depicted in orange and blue in Fig. 9. The red dotted line represents the curve of a chance predictor. Both the orange and blue curve are notably separated from the red curve. However, the orange curve of the proposed method with feature fusion shows a higher true positive rate at nearly all false positive rates when compared to the blue curve of the LSTM. This is further supported by their average AUC scores of 80.60% and 71.88%, respectively.

**Figure 9:**
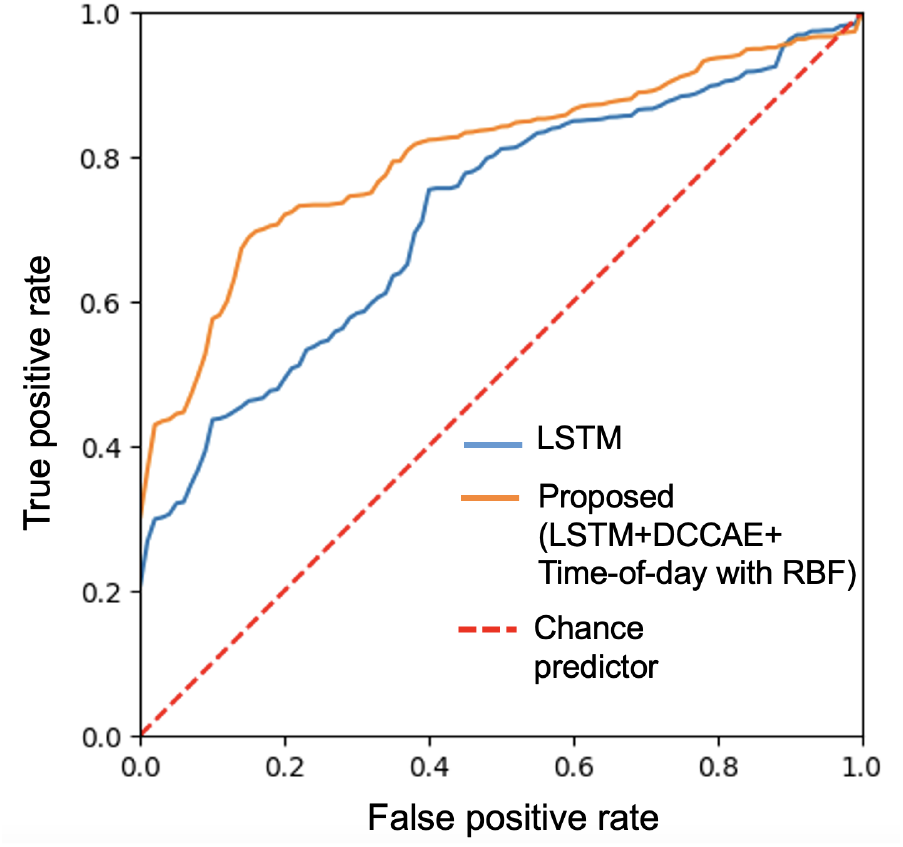
ROC for the proposed method fusing features from LSTM, DCCAE, and time-of-day using RBF encoding, along with the baseline LSTM. The red dotted line denotes the ROC of the chance predictor.

### 4.3 Growing neural network for feature fusion

We also compared the proposed growing network approach with the typically employed all-at-once approach, where we train a new neural network for classification from the beginning instead of letting a pre-trained network grow incrementally. The 10-fold cross-validation results averaged over all folds, are presented in Table 2. The proposed method fusing all three features with growing network achieves higher accuracy compared to the all-at-once approach. Compared to the baseline LSTM, the average accuracy for the all-at-once approach is even lower. However, this does not mean that the all-at-once approach always performs worse than the baseline. We present the accuracies for each fold in Figure 10, sorted in increasing order of the baseline LSTM accuracy. The proposed growing network approach outperforms the baseline in folds with comparably low accuracy (indicated by the green-shaded region) while performing equally well in folds with high baseline accuracies. In contrast, the all-at-once approach performs worse than the baseline in seven out of ten folds, while it outperforms the baseline in three cases, resulting in lower average accuracy as shown in Table 2. Moreover, the all-at-once approach performs worse in folds where the baseline already achieves high accuracy, as indicated by the red-shaded regions.

**Table 2:** Performance of the proposed method for feature fusion with all-at-once and growing network.

| Method | Accuracy |
| --- | --- |
| LSTM | 74.44% |
| Proposed (all-at-once) | 72.24% |
| Proposed (growing) | **81.70%** |

**Figure 10:**
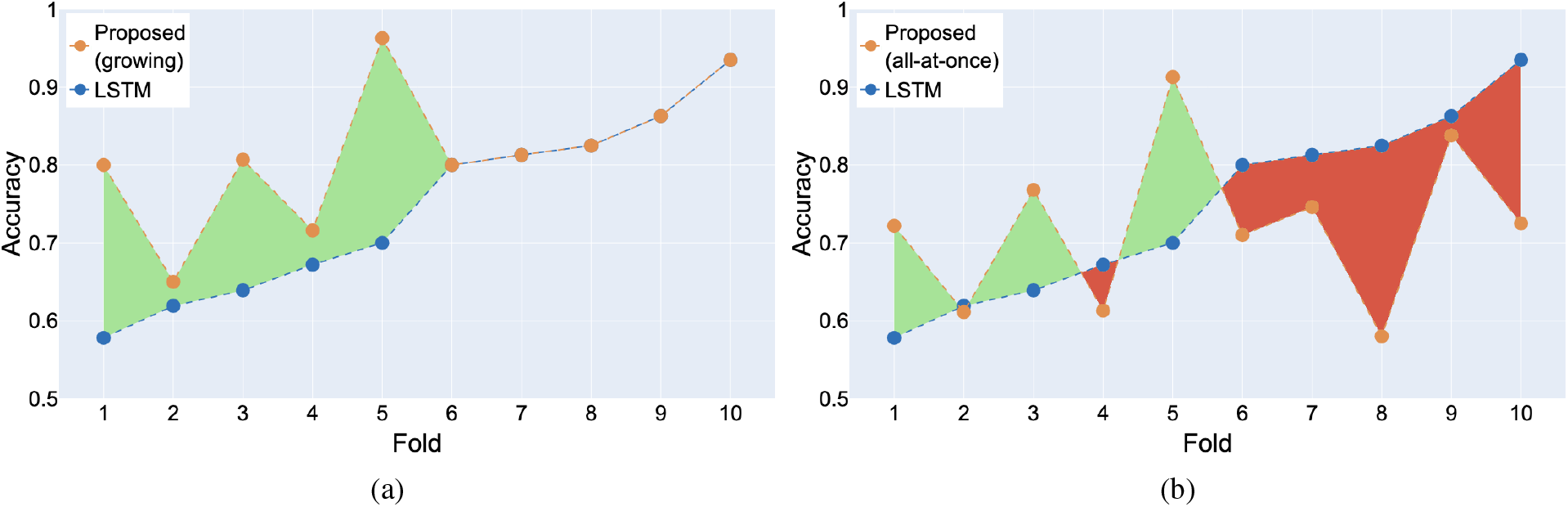
Prediction accuracy per fold for the proposed method of feature fusion using a) the growing network and b) the all-at-once approach is compared to the LSTM. The improvement in performance is shown in the green-shaded region, while the decrease in performance is indicated by the red-shaded region.

### 4.4 Prediction performance for individual and subgroups of patients based on clinical data

The prediction accuracy of the proposed method for each patient and the baseline LSTM using leave-one-patient-out cross-validation is shown in Fig. 11a. The proposed method achieves better-than-chance accuracy for 97.3% of the patients, while the LSTM achieves it for 84.2% of the patients. Averaged across all patients, the proposed method achieves 73.42% accuracy, outperforming the baseline LSTM approach, which attains 64.15%. Moreover, for 84% of the patients, the proposed method improves accuracy over the LSTM. This is shown in Fig. 11b, which illustrates the patient-wise improvement from low to high. We can see that for three patients where accuracies decreased, the difference in accuracy is minimal, and for three other patients, the accuracy is the same for both LSTM and the proposed method. This indicates that the performance improvements generalize well on a patient level.

**Figure 11:**
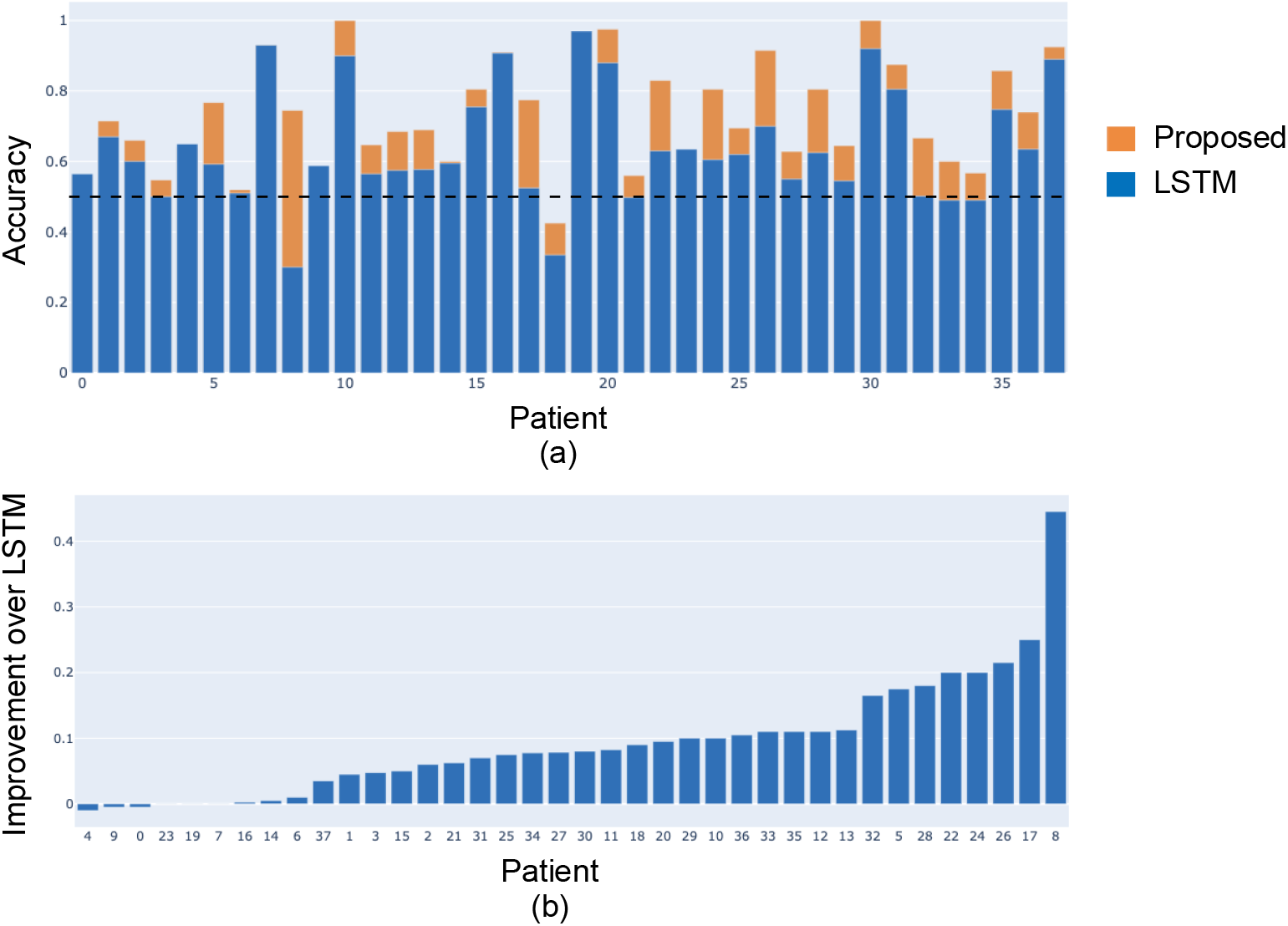
Performance of the proposed method at the patient level using leave-one-patient-out evaluation. a) Accuracy for different patients. The black dotted line indicates the accuracy of a chance classifier. b) Improvement of the proposed method over the baseline LSTM for each patient.

We post-hoc analyzed the differences in prediction performance across patients grouped based on their clinical data. The results are presented in Fig. 12. The proposed method performs slightly better for patients with GTC seizures than for patients with FBTC seizures (Fig. 12a). The performance for males and females is nearly identical (Fig. 12a). Due to the low number of patients with genetic, infectious, and immune etiology types, we grouped them into a single subgroup for this analysis. The method’s accuracy is slightly higher for patients with structural etiology and even higher for patients with unknown etiology compared to those with other etiology types (Fig. 12c). For patients with a seizure frequency of less than 15 seizures per month within 30 days of their LTM admission, the prediction performance is better than for patients with a seizure frequency greater than 15 (Fig. 12d). We also analyzed the accuracy of the method with respect to the epilepsy duration computed as the difference between the age of enrollment and the age of the first seizure. We observe that, on average, the method has higher prediction accuracy for patients with larger epilepsy duration. We did not observe significant differences between the accuracy of different subgroups within each clinical variable. To analyze the relationship between the accuracy and the combination of various clinical variables, we computed canonical correlation between the accuracy and the combination of etiology, seizure frequency, and epilepsy duration. We observed a significant canonical correlation coefficient of 0.51 (p-value = 0.016).

**Figure 12:**
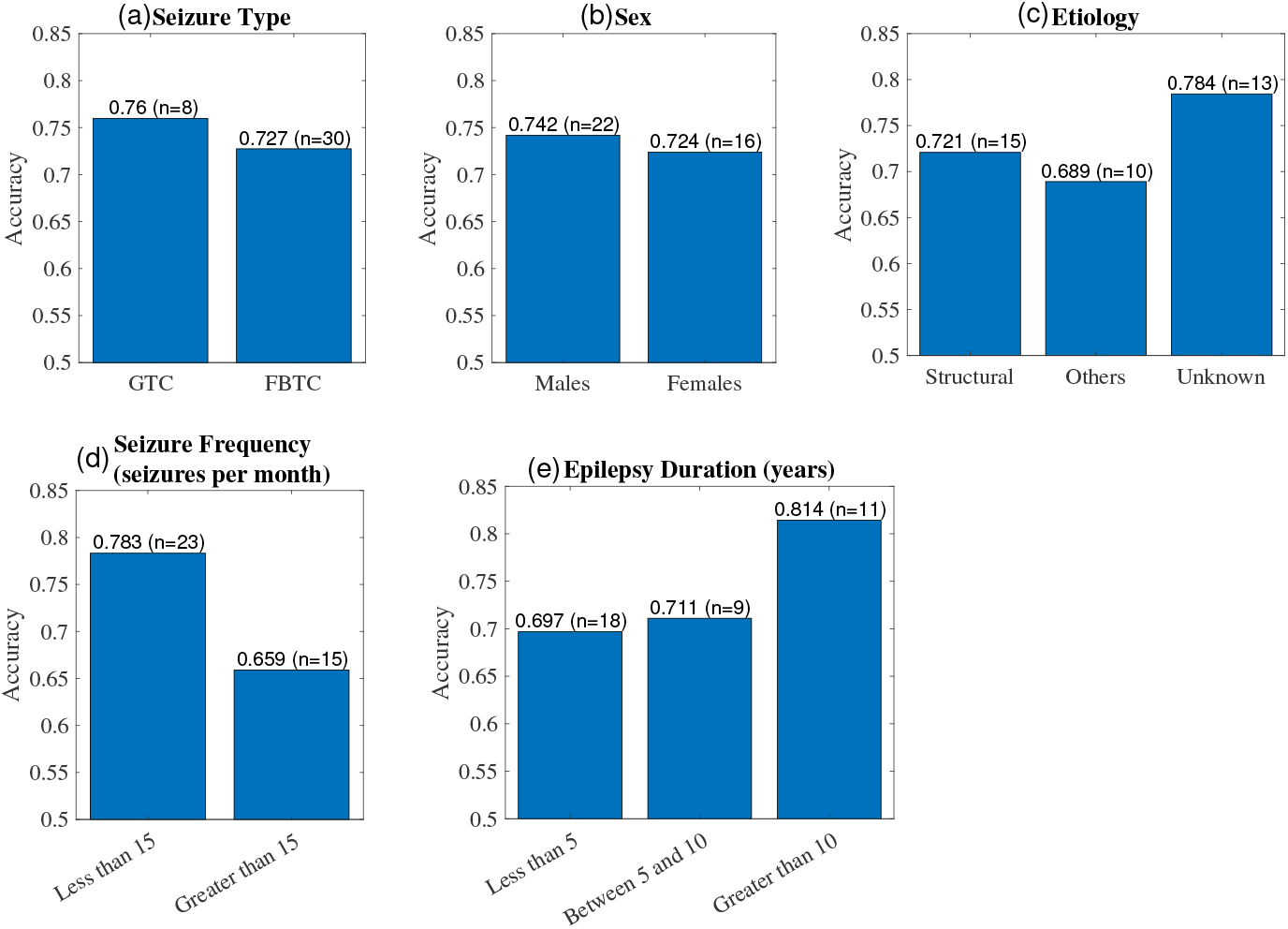
Prediction accuracy for patient subgroups based on their clinical data. a) Seizure type, b) sex, c) etiology, d) seizure frequency, and e) epilepsy duration. A significant canonical correlation of 0.51 (p-value = 0.016) is obtained between accuracy and the combination of etiology, seizure frequency, and epilepsy duration.

### 4.5 Effect of data quality on prediction performance

To evaluate the performance of the proposed method with the data quality of patient recordings, we varied the threshold for the data quality score used in the inclusion criteria. The data quality threshold was adjusted between 0.85 and 0.5. Testing below 0.5 was not feasible, as all patients had a data quality score exceeding 0.5. Testing above 0.85 was avoided due to the very small number of patient data available at higher thresholds. The average accuracy of the proposed method across varying data quality thresholds is shown in Table 3. The higher-quality data with thresholds of 0.75 and 0.85 provide higher performance, whereas the inclusion of lower-quality data degrades performance, even though more patient data becomes available at lower thresholds.

**Table 3:**
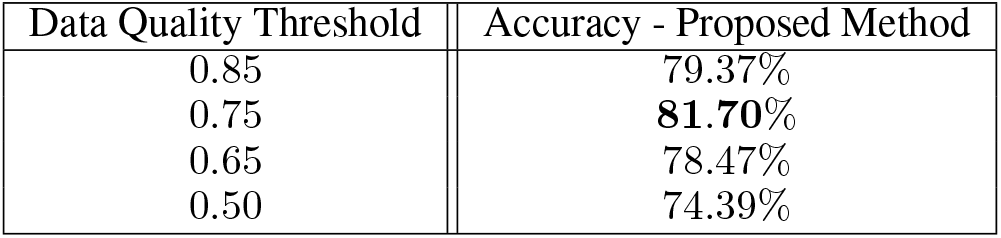
Evaluation of the proposed method for varying wearable data quality threshold.

### 4.6 Effect of preictal data duration on prediction performance

We also evaluated the method’s performance on data that is temporally close to or distant from the seizure onset by varying the total duration of preictal data used for training the model. Table 4 presents preictal data durations ranging from 2 to 30 minutes. The highest accuracy is achieved with a 5-minute duration, with a slight decrease in performance observed for 3-minute and 2-minute durations. As we include data further from the seizure onset, the accuracy decreases, reaching its lowest when using preictal data with a 30-minute duration.

**Table 4:**
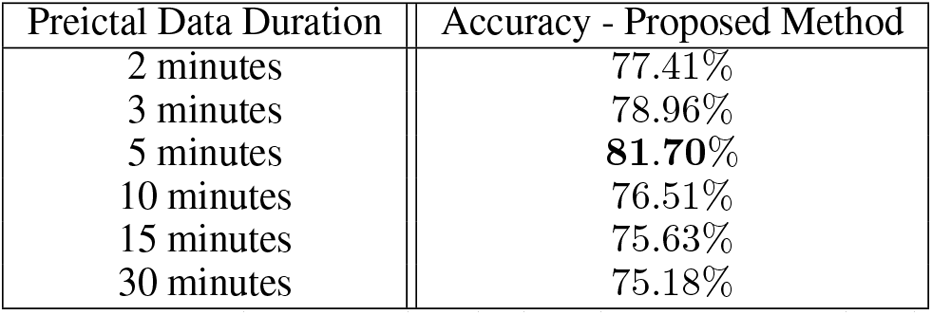
Evaluation of the proposed method for different preictal data duration.

| Preictal Data Duration | Accuracy - Proposed Method |
| --- | --- |
| 2 minutes | 77.41% |
| 3 minutes | 78.96% |
| 5 minutes | **81.70%** |
| 10 minutes | 76.51% |
| 15 minutes | 75.63% |
| 30 minutes | 75.18% |

## 5 Discussion

We present a patient-agnostic seizure prediction method for tonic-clonic seizures. Our novel approach fuses features from: a supervised LSTM model, an unsupervised correlation-maximizing DCCAE, and time-of-day features computed using trigonometric and RBF encodings. These features are fused using a growing neural network. The highest accuracy achieved is 81.7%, obtained by fusing all three feature sets with RBF encoding using 10-fold cross-validation. Therefore, we may be able to predict seizures from the wearable data with incremental accuracy based on timing and input features [22, 23, 24, 25]. Our method captures patient-independent patterns in the data, showing feasibility in real-world scenarios, as it does not require extensive seizure data from new patients potentially predicting from the first seizure. Such a model is easier to fine-tune, requiring only a small amount of patient-specific data for adjustment rather than retraining a new model for each patient. This provides a major advantage over many existing seizure prediction methods that train patient-specific models, requiring a large amount of labeled patient-specific data [9, 26, 13].

## 5.1 Feature fusion from different techniques leads to improved prediction performance

In our seizure prediction method, we computed features using three different techniques. The supervised LSTM extracts features from the time series segments of EDA, HR, and TEMP, aiming to distinguish between preictal and interictal segments maximally. Consistent with other studies on wearable-based seizure prediction using LSTM models [23, 26, 24], this approach achieves a prediction accuracy of 74.44%. However, the LSTM focuses on maximizing classification accuracy without explicitly accounting for the interactions between different ANS modalities and the changes in those interactions before the seizure onset.

In contrast, DCCAE learns features that capture nonlinear relationships between different ANS subsystems measured by EDA and HR. Thus, DCCAE could reduce the high intra- and interpatient variability inherent to the ANS by focusing on multimodal interactions. We have shown that canonical correlations extract multimodal interactions between the ANS subsystems [54, 55], and the mean canonical correlation across all modalities changes as a seizure nears, offering potential prediction biomarker [31, 37]. Fusing DCCAE features with those from the LSTM increased prediction accuracy by 2.21% to an average accuracy of 76.65%, indicating that DCCAE provides complementary information to the LSTM.

The third set of features we incorporated relates to the 24-hour patterns, as seizures are known to follow cyclic patterns [38, 39]. Due to the limited data available from the inpatient setting, we included patient-agnostic time-of-day information from the corresponding preictal and interictal segments, using both trigonometric (sine and cosine encodings) and RBF encodings. Both encodings enhanced prediction performance, with trigonometric encoding improving accuracy by 1.68% and RBF encoding leading to a 5.05% improvement over the fused LSTM and DCCAE features. Our results align with the results from [41, 42, 40] and show that including 24-hour patterns improves prediction performance. Compared to [41, 42], which included patient-specific seizure-occurrence patterns, we show that the performance improvement can be translated to a patient-agnostic prediction model similar to [40]. Compared to [40], which also included trigonometric encoding combined with wearable data as a single input to the neural network, we show that RBF encoding leads to higher performance, and the performance is further improved by fusing them using a growing neural network. We use 24 RBF functions to explicitly represent each hour of the day so that the network can directly capture hourly patterns in the data. Thus, RBF encoding is an interpretable method for incorporating cyclic seizure-related features with the potential for extension to model multiday and monthly patterns.

The trend of features from the DCCAE and the time-of-day improving the accuracy of the baseline LSTM model is similar on the individual patient level using leave-one-patient-out evaluation, indicating the complementary information these features could contain for seizure prediction. For 32 out of 38 patients, the accuracy of the proposed method is higher than the baseline LSTM. The accuracy is unchanged for three patients, while there is a marginal decrease in accuracy for the remaining three patients. This could be due to an increased model complexity due to the additional features compared to the complementary predictive information they contain for these three patients. Our post-hoc analysis showed that our method’s performance varies with patients’ clinical characteristics. We observed differences in the average accuracy across patient subgroups based on etiology, seizure frequency, and duration of epilepsy and a significant canonical correlation coefficient between accuracy and the combination of etiology, seizure frequency, and epilepsy duration. Therefore, differences in clinical characteristics could lead to differences in prediction performance, and patients’ clinical characteristics should be considered when designing seizure prediction and forecasting methods.

## 5.2 Growing neural network is better suited for combining multimodal heterogeneous data than an all-at-once network

The three feature sets in this study are extracted using different techniques, and the strategy used to combine these features can significantly influence the final prediction performance. We adopted a growing neural network methodology that incrementally expands the network size during training. Our results indicate that the growing network outperforms the traditional all-at-once strategy by an average accuracy difference of 9.5%. In some evaluation folds, where the LSTM achieves comparatively low prediction accuracy, the all-at-once strategy performs similarly to the growing network and enhances the prediction performance by fusing different features. However, it performs worse in evaluation folds where the LSTM achieves relatively high accuracies. In contrast, the growing strategy is designed to perform equally as well as or better than the baseline LSTM for all folds. The growing neural network also benefits from initializing its weights using the LSTM, which improves the extraction of predictive information from different features. The advantage of initializing weights for seizure prediction has also been shown using transfer learning with EEG data [56].

## 5.3 Wearable data quality affects prediction performance

Epileptic seizures impact the quality of data collected from wearable devices, as shown in [57], where tonic-clonic and motor movements during seizures reduce the wearable data quality, especially BVP. However, no study has yet assessed the influence of wearable data quality on seizure prediction accuracy. In this study, we evaluated data quality during the interictal and preictal periods for each patient across three modalities of EDA, HR (using BVP), and TEMP. We showed that including lower-quality data reduces prediction accuracy with the lowest-quality threshold, resulting in the lowest prediction accuracy. This occurred despite the availability of additional training data at lower thresholds, indicating that wearable data quality impacts the prediction performance. These results emphasize the importance of data quality in seizure monitoring, similar to [44, 57, 58], and indicate that data quality should be a critical consideration in wearable-based seizure prediction. Particularly for the outpatient setting, if the input data quality falls below a certain threshold, the data may either be pre-processed, or the method could skip the prediction for that data segment to ensure reliable seizure prediction.

## 5.4 Preictal data duration impacts prediction accuracy

Preictal autonomic changes exhibit high variability across patients and seizures, and the time at which these changes can be detected before seizure onset also varies, even within the same patient [59, 60]. The length of preictal data used in different seizure prediction methods is not standardized. For example, [26] and [23] used one hour as the preictal data duration, whereas [19] used 15 minutes. In this study, we analyzed the impact of preictal data duration on the prediction performance of our proposed method. A 5-minute duration yielded the best accuracy, with the performance decreasing slightly when the duration was reduced to 3 minutes and 2 minutes. This decrease could be due to the reduction in the available training data or indicate that the autonomic changes occurring 5 minutes before seizure onset are most discriminatory for our method. Conversely, the prediction accuracy decreased significantly as the preictal data duration increased, with the worst performance observed at 30 minutes. This finding aligns with our previous results, where we showed that multimodal autonomic interaction increased significantly before seizure onset but were at baseline level around 30 minutes or more prior to the seizure [31]. An improved approach would involve determining an optimal preictal duration for each patient, as proposed in [61]. However, this approach would need longitudinal data that includes numerous seizures per patient along with corresponding preictal data instances, which is not feasible with our dataset.

## 5.5 Limitations and challenges

Our proposed method has limitations in both the methodological and clinical aspects.

### 5.5.1 Methodological limitations

For DCCAE, we focussed on HR and EDA, as seizure-related changes have been detected in peripheral recordings using these modalities [62, 63, 30, 64]. The DCCAE could, however, be extended by incorporating additional autonomic signals such as the TEMP or respiratory rate [31]. [23] showed that the seizure prediction performance in a pseudo-prospective set-up improved when all wearable modalities were included but worked in 43% of the patients. This indicates preictal changes across all modalities. Extending our method to more than two modalities would also necessitate a change in the cost function of DCCAE using multiset CCA and taking different correlation structures into account [65, 66]. This extension would increase the network complexity and the number of training parameters. With the limited data from tonic-clonic seizures, we chose EDA and HR for this study, and the extension of adding correlations across more modalities is left for our future work with a larger cohort of patients with more seizure types. Additionally, with the larger data set, more complex neural network architectures, such as attention models, can be incorporated [67]. To address overfitting in LSTM and DCCAE, we used L2 regularization. Future work, including attention models, would require advanced regularization techniques for tackling overfitting such as dropout and complexity regularization [68, 69, 70].

### 5.5.2 Clinical limitations

From the clinical perspective, the main limitations of this work are patient selection, recording setup, and the inclusion of only autonomic recordings. Our cohort includes children admitted to the LTM. Although the included patients represent a pediatric population spanning all age groups from one to 23 years, with an approximately uniform distribution of seizure onset times, all patients were referred by their clinicians to the LTM specialized in diagnosing and treating epilepsy. This introduces a selection bias by including only patients with a particular disease severity. Recording the data in the LTM allows for gold-standard seizure characterization. However, it also limits the transferability of our results to the outpatient setting, an environment under less control.

Furthermore, the patients may have been weaned off the anti-seizure medications during the monitoring and may have imaging and EEG findings related to seizures, and thus, the confounding effects cannot be ruled out. We analyzed tonic-clonic seizures, excluding other seizure types. While this selection approach enhances the homogeneity in our dataset, it also limits our dataset’s size and reduces our findings’ generalizability to other seizure types. Another limitation of our method is the inclusion of only autonomic data for our prediction method and the absence of additional clinical information such as seizure frequency and anti-seizure medication reduction, which have been shown to play a role in seizure prediction [71, 72]. In the context of the limited dataset, including clinical variables could bias the method and, thus, necessitates careful cohort generation with a balanced distribution of clinical variables. This would have further reduced the size of our dataset. Hence, we opted for a post-hoc analysis showing patients’ accuracy in subgroups with different clinical characteristics. As part of our future work, we aim to integrate the clinical variables as input to our method using a larger dataset that includes different seizure types.

## 6 Acknowledgments

The study was supported by the Paderborn University Research Award and partially by the Epilepsy Research Fund.

